# A distinctive PI(4,5)P_2_ compartment forms during entosis and related engulfment processes

**DOI:** 10.1101/2024.10.22.619613

**Authors:** Joanne Durgan, Katherine Sloan, Marie-Charlotte Domart, Lucy M Collinson, Oliver Florey

## Abstract

Entosis is a form of cell cannibalism prevalent in human tumours. During entosis, one live and viable epithelial cell is completely internalised by another, then housed inside a large, single-membrane, endolysosomal vacuole in the host cytosol. The composition and maturation of this specialised, macroendocytic compartment has not been fully defined, but ultimately, the inner cell is killed and digested by host lysosomes. Here, we investigate the molecular characteristics and maturation profile of the entotic vacuole. Like phagosomes and macropinosomes, this vacuole undergoes a series of phospholipid modifications, but its maturation profile bears distinctive dynamics. While PI(4,5)P_2_ is lost rapidly during phagosome maturation, entosis yields an unusual, intracellular PI(4,5)P_2_-positive compartment, that can persist for hours, suggesting vacuole maturation is uncoupled from membrane scission. Loss of PI(4,5)P_2_ is eventually triggered, a requisite step for lysosomal killing of the internalised cell. More broadly, PI(4,5)P_2_-positive vacuoles also form during T-cell engulfment by thymic nurse cells, dependent on ROCK activity, suggesting this distinctive compartment represents a shared feature of entosis-like cell engulfments.

## Introduction

During macroendocytic processes, such as macropinocytosis and phagocytosis, cells engulf external materials, including fluids, pathogens or apoptotic debris (Elliott and Ravichandran, 2010; Swanson and King, 2019). These engulfment events play fundamental roles in diverse processes, including immune defence, tissue remodelling and nutrient uptake (Lim et al., 2017). Mechanistically, cell engulfment involves cargo binding and co-ordinated engagement of the cytoskeleton, to extend the plasma membrane and envelop the target (Mylvaganam et al., 2021). The cargo is then internalised into a vacuole compartment, derived from invagination of the plasma membrane, which undergoes a complex series of maturation steps, including sequential phosphoinositide (PI) lipid changes and protein recruitments, driving a transition from early to late endosome/lysosome identity (Levin et al., 2015). These changes include loss of PI(4,5)P_2_, generation of PI3P, recruitment of Rab5 and 7, generation of PI(3,5)P_2_ and lysosome fusion (Desjardins et al., 1994); ATG8/LC3 decoration can also occur through non-canonical autophagy (Florey et al., 2011). Proper vacuole maturation, and lysosome fusion, are essential in determining the fate of the internalised cargo.

Cell-in-cell formation is a specialised form of engulfment, involving internalisation of one live cell by another. It occurs in a range of contexts, often followed by lysosomal killing of the engulfed cell by its host (Li et al., 2015; Sterling et al., 2023). Heterotypic engulfments include thymocyte internalisation by thymic nurse cells (Philp et al., 1993), and immune cell engulfment by hepatocytes or cancer cells (Davies et al., 2019). Homotypic cell cannibalism occurs in epithelial cells through entosis, driven by actomyosin contractility in the internalised cell, which ‘invades’ into its host (Overholtzer et al., 2007; Sun et al., 2014). This can be induced by loss of matrix adhesion, dysregulated mitosis and low glucose (Durgan et al., 2017; Hamann et al., 2017; Overholtzer et al., 2007), all hallmark features of cancer (Durgan and Florey, 2018). Entosis is prevalent in human tumours, associated with higher tumor grade and poor prognosis (Fais and Overholtzer, 2018; Mackay and Muller, 2019; Ruan et al., 2019; Song et al., 2023), and mediates complex effects on tumour promotion (Krajcovic et al., 2011), suppression (Overholtzer et al., 2007) and evolution (Sun et al., 2014).

During entosis, the internalised cell is housed inside a single-membrane vacuole in the host cytosol, where it can remain viable for several hours. Like phagosomes, this vacuole can recruit ATG8s, and the inner cell is ultimately killed and digested by host lysosomes (Florey et al., 2011). However, little more is known about the composition and processing of this specialised, macroendocytic compartment. In this study, we examined the maturation profile of the entotic vacuole, alongside phagocytosis and T-cell/thymic nurse cell engulfment. We report the identification of a distinctive PI(4,5)P_2_-positive, intracellular compartment formed during entosis-like engulfment processes, and show that loss of PI(4,5)P_2_ is required for inner cell killing.

## Results and Discussion

### Differential actin and phosphoinositide dynamics during phagocytic and entotic engulfment

To investigate the molecular characteristics of entotic vacuole maturation, a comparative analysis with phagocytosis was performed, with an initial focus on actin and phosphoinositide dynamics. Mouse macrophage (J774.1A) were used to model phagocytosis, alongside human epithelial cells that undergo entosis upon matrix deadhesion (MCF10A). Fluorescent labels, and genetically encoded biosensors, were used to visualise actin, phosphatidylinositol (3,4,5)-trisphosphate (PI(3,4,5)P_3_; Akt-PH-GFP) and phosphatidylinositol (4,5)-bisphosphate (PI(4,5)P_2_; GFP-PLCδ-PH). For entosis, cell pairs were selected in which only the outer cell expressed detectable fluorescent biosensors, to focus exclusively on the entotic vacuole signal.

During phagocytosis, actin and PI(3,4,5)P_3_ enrich strongly at the phagocytic cup (Fig. 1A-B), consistent with previous reports (Freeman and Grinstein, 2014; Henry et al., 2004; Marshall et al., 2001). A transient accumulation of PI(4,5)P_2_ is also observed, throughout the plasmalemma and phagocytic cup, which then disappears from the base and is completely lost upon closure and completion of the phagosome (Fig. 1C, Movie 1) (Botelho et al., 2000; Scott et al., 2005).

**Figure 1.**
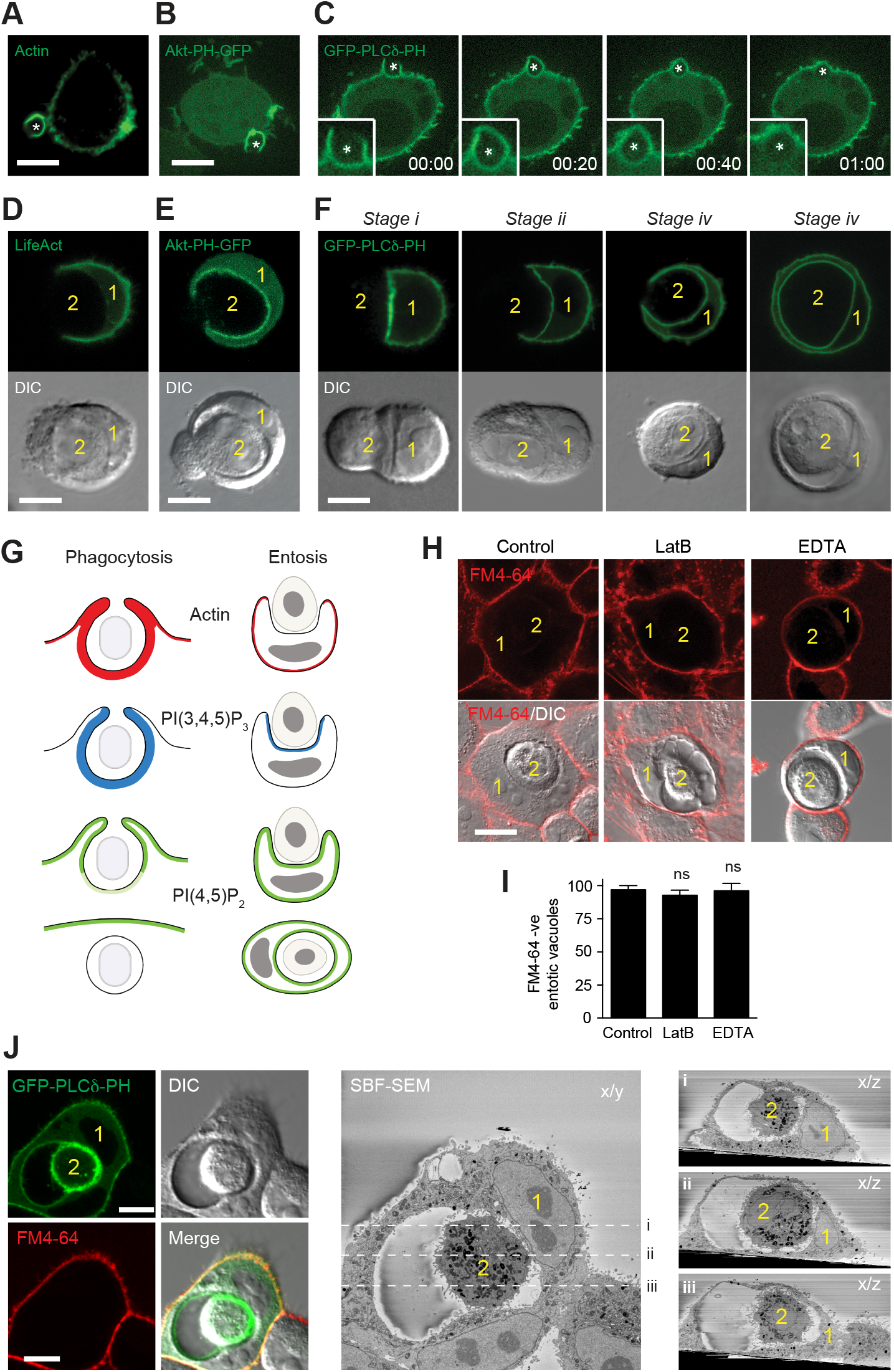
Phagosome versus entotic vacuole maturation. (A-C) Confocal images of J774.1A cells incubated with opsonised zymosan, * = zymosan particle. (A) Stained for actin, with phalloidin (B) Expressing Akt-PH-GFP, PIP3 sensor. Scale bars, 5 μm. (C) Live cells expressing GFP-PLCδ-PH, PIP2 sensor. Zoomed inset of forming phagosome. Time, min:sec. (D-F) Live confocal images of MCF10A cells grown in soft-agar. Fluorescent and DIC images show forming entotic structures, with outer (1) and inner (2) cells indicated, and outer cells expressing (D) GFP-LifeAct, to mark actin, (E) Akt-PH-GFP, to mark PIP3 (F) GFP-PLCδ-PH, to mark PIP2; representative images show progressive stages of engulfment (*i-iv*). Scale bar, 10 μm. (G) Schematic comparison of actin and phosphoinositide dynamics during phagocytosis and entosis. (H) Confocal images of MCF10A entotic structures, labelled with FM4-64, -/+ Latrunculin B, 1 μM or EDTA, 5 mM pre-treatment. Outer cell (1), inner cell (2). Scale bar, 15 μm. (I) Quantification of (H), data represent mean±SEM from 3 independent experiments (n>50). (J) 3D CLEM analysis of MCF10A entosis; outer cell (1), inner cell (2). Live cell confocal microscopy images of DIC, GFP-PLCδ-PH (PIP2, in both cells) and FM4-64, with SBF-SEM x/y axis, (*i-iii*) show x/z SBF-SEM slices from within the entotic structure. Scale bar, 15 μm.

During entosis, lower levels of actin are observed at the entotic cup, relative to the plasma membrane (Fig. 1D), contrasting with phagocytosis. Nevertheless, PI(3,4,5)P_3_ enriches at the cell-cell interface (Fig. 1E), consistent with its generation at the cell junctions which drive entotic engulfment. PI(4,5)P_2_ is clearly observed at both the plasmalemma and entotic cup (Fig. 1F). Strikingly, and somewhat unexpectedly, this PI(4,5)P_2_ persists through all stages of entotic engulfment (i-iii), and beyond (iv), suggesting this species is not lost following scission during entosis, as it is in phagocytosis.

Together, these data reveal both similarities and key differences in actin and phosphoinositide dynamics, during different cell engulfment processes (Fig. 1G). These distinctions reflect different mechanisms of engulfment, where phagocytosis is mediated by the ‘outer’ cell, while entosis is largely driven by the ‘inner’ cell, which actively invades into its host, based on biophysical differences in RhoA/ROCK mediated contractility (Overholtzer et al., 2007; Sun et al., 2014). Loss of actin at the entotic cup, in the more deformable host cell, may permit its invasion by a stiffer, more contractile neighbour. Alongside this difference in actin dynamics, entosis also displays a distinctive lipid profile, yielding an unusual, PI(4,5)P_2_-positive compartment inside the host cell.

### PI(4,5)P_2_ positive entotic vacuoles have undergone complete scission

The apparent persistence of PI(4,5)P_2_ on entotic vacuoles could potentially be explained by incomplete engulfment/scission from the plasma membrane. To exclude this, a lipophilic dye (FM4-64) was added to entotic cell pairs, to stain any membrane exposed to media.

Strikingly, while FM4-64 clearly labels the host cell plasma membrane, entotic vacuoles and inner cells remain FM4-64 negative (Fig. 1H), consistent with complete internalisation. We questioned whether tight junctions might complicate this interpretation by impeding FM4-64 diffusion in a partially engulfed structure, but treatment with latrunculin B to target actin (Shen and Turner, 2005), or EDTA to chelate calcium (Ye et al., 1999), had no effect (Fig. 1H-I). To further reinforce the conclusion that PI(4,5)P_2_ -positive, FM4-64 negative entotic cells are completely internalised, ultrastructural analyses were performed by 3D Correlative Light and Electron Microscopy (CLEM). Cell-in-cell pairs were imaged by live confocal microscopy, then processed for Serial Block Face Scanning Electron Microscopy (SBF-SEM), to visualise the entire volume of an entotic structure (Fig. 1J, Movie 2). Careful analysis of multiple samples confirmed that each inner cell appeared completely internalised, with the vacuole membrane surrounding it in full, and no evidence of cellular protrusions extending outside of the host cell, in any slice of the full 3D structure. In contrast, 3D CLEM analysis of FM4-64 positive structures revealed obvious structural evidence of incomplete engulfment, further strengthening our interpretation (Fig. S1).

Together, these data confirm that following complete scission from the plasma membrane, the entotic vacuole remains positive for PI(4,5)P_2_, a species typically associated with plasma membrane identity (Wills and Hammond, 2022) for an extended time, often several hours. This contrasts with phagosomes, which rapidly lose PI(4,5)P_2_ upon formation and maturation to a late endosome/lysosome state, suggesting this composition is a distinctive feature of this specialised intracellular vacuole.

### Formation of a PI(4,5)P_2_-positive vacuole following T-cell internalisation into thymic nurse cells

Alongside entosis, a variety of cell-in-cell formation processes occur in different cell types, mediated by different engulfment processes (He et al., 2013). For instance, live T-cells and thymocytes can invade into specialised epithelial thymic nurse cells (TNCs) (Guyden et al., 2015; Li et al., 1992; Philp et al., 1993). To extend this study, we explored the vacuolar characteristics associated with this alternative form of cell engulfment. A TNC line, TNCR3.1 (Nishimura et al., 1990), stably expressing the PI(4,5)P_2_ reporter (GFP-PLCδ-PH), was incubated with T-cell hybridoma cells, DO11.10, pre-labelled with Hoechst. Using confocal microscopy, different stages of their interaction were captured; a representative example is presented (Fig. 2A-B). Initially, the DO11.10 T-cell overlays the adherent, TNCR3.1 cell (Stage 1). During early engagement (Stage 2), the T-cell migrates beneath the TNC and pushes up into it, effectively invading from below. By Stage 3, the T-cell is almost fully internalised, with only a small aperture remaining at the basal side of its TNC host. Finally, the T-cell appears to be fully internalised, housed inside a vacuole in the TNC cytosol, which remains PI(4,5)P_2_-positive (Stage 4).

**Figure 2.**
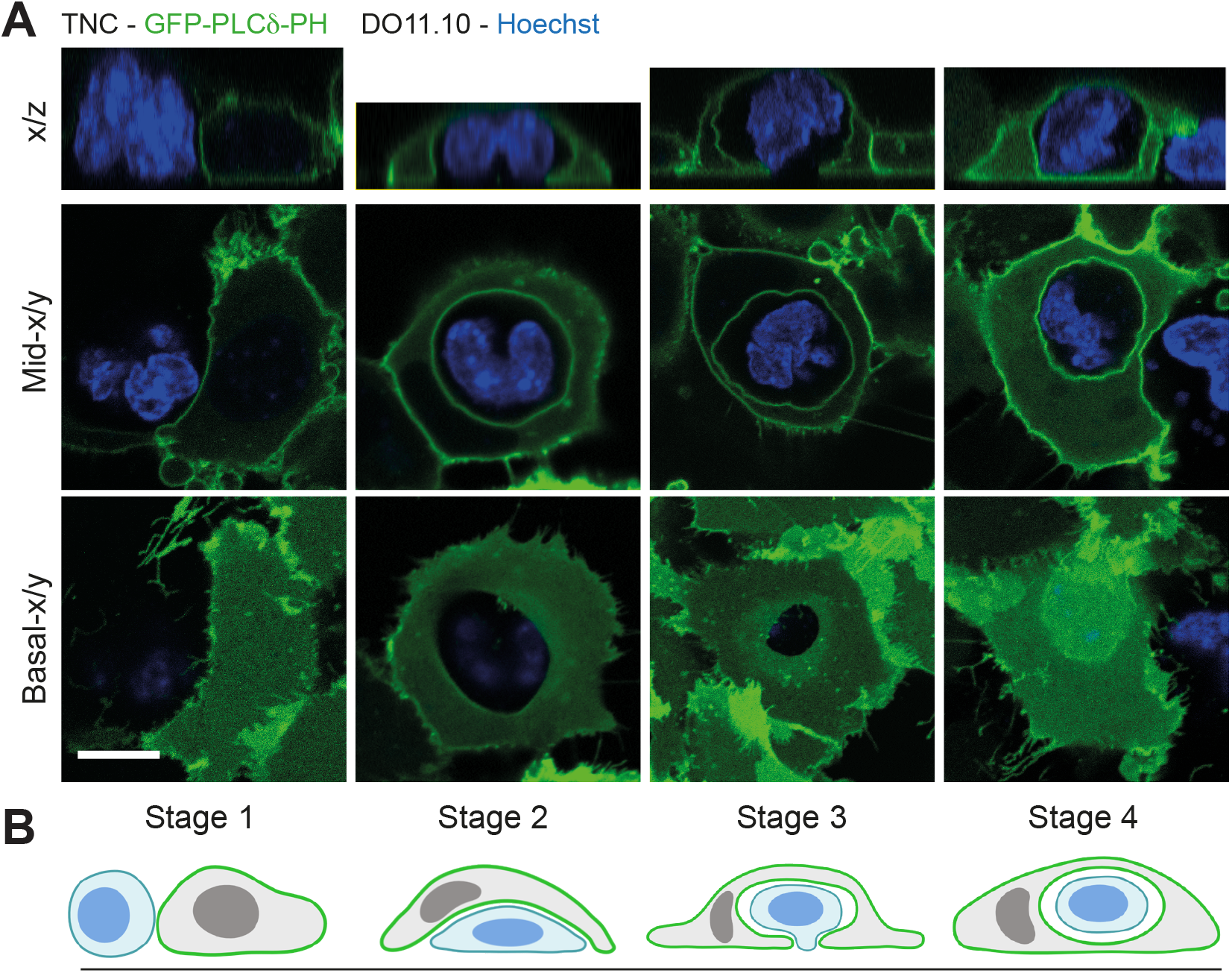
PI(4,5)P2 dynamics during thymic nurse cell/T-cell engulfment. (A) Live confocal images of TNC3R.1 cells expressing GFP-PLCδ-PH (PIP2, green), co-cultured with Hoechst-labelled DO11.10 cells (DNA, blue). Images show basal and mid x/y slices from confocal stacks and x/z images, through progressive stages of cell-cell interaction (1-4). Scale bar, 10 μm. (B) Schematic representation of the different stages of TNCR3.1 (grey) and DO11.10 (blue) cell interactions/engulfment; PI(4,5)P2 (green).

These data reveal that, like entosis, the engulfment of T-cells by TNCs yields an intracellular, PI(4,5)P_2_-positive vacuole, indicating this may represent a broader, shared feature among certain engulfment events.

### Formation of PI(4,5)P_2_ positive compartments is dependent on the mode of engulfment, not target cell viability

Entosis and TNC/T-cell internalisation involve live cell engulfment events, while phagocytosis classically targets dead or dying cells. These engulfment processes also proceed through different mechanisms, driven by either host (phagocytosis) or prey (entosis). We next questioned whether formation of a PI(4,5)P_2_-positive compartment is dependent on the viability of the internalised cell, or the mechanism of its engulfment.

To distinguish between these possibilities, overlapping models were compared: i) entosis-like engulfment of live target cells (TNCR3.1, viable DO11.10 cells), ii) phagocytic uptake of dead target cells (TNCR3.1, apoptotic DO11.10 cells), iii) phagocytic uptake of live target cells (J774.1A, viable Jurkat coated in anti-CD47 (Métayer et al., 2017), iv) phagocytic uptake of dead target cells (J774.1A, apoptotic Jurkat). In each experiment (Fig. 3), the host cells expressed PI(4,5)P_2_ reporter, while inner cells were pre-stained for DNA; viability was confirmed by nuclear morphology, with apoptotic cells displaying characteristic concentrated and fragmented staining. Cells were also treated with FM4-64 to confirm completion of engulfment. Consistent with Fig. 2, PI(4,5)P_2_-positive, FM4-64 negative vacuoles were detected following the entosis-like internalisation of live T-cells in TNCs (Fig. 3A and E). In contrast, no PI(4,5)P_2_-positive, FM4-64 negative compartments were observed in any of the phagocytic conditions, regardless of whether the target cell was dead or alive (Fig. 3B-E). These data indicate that formation of PI(4,5)P_2_-positive intracellular vacuoles is a distinctive feature of entosis-like engulfments, rather than live cell internalisations.

**Figure 3.**
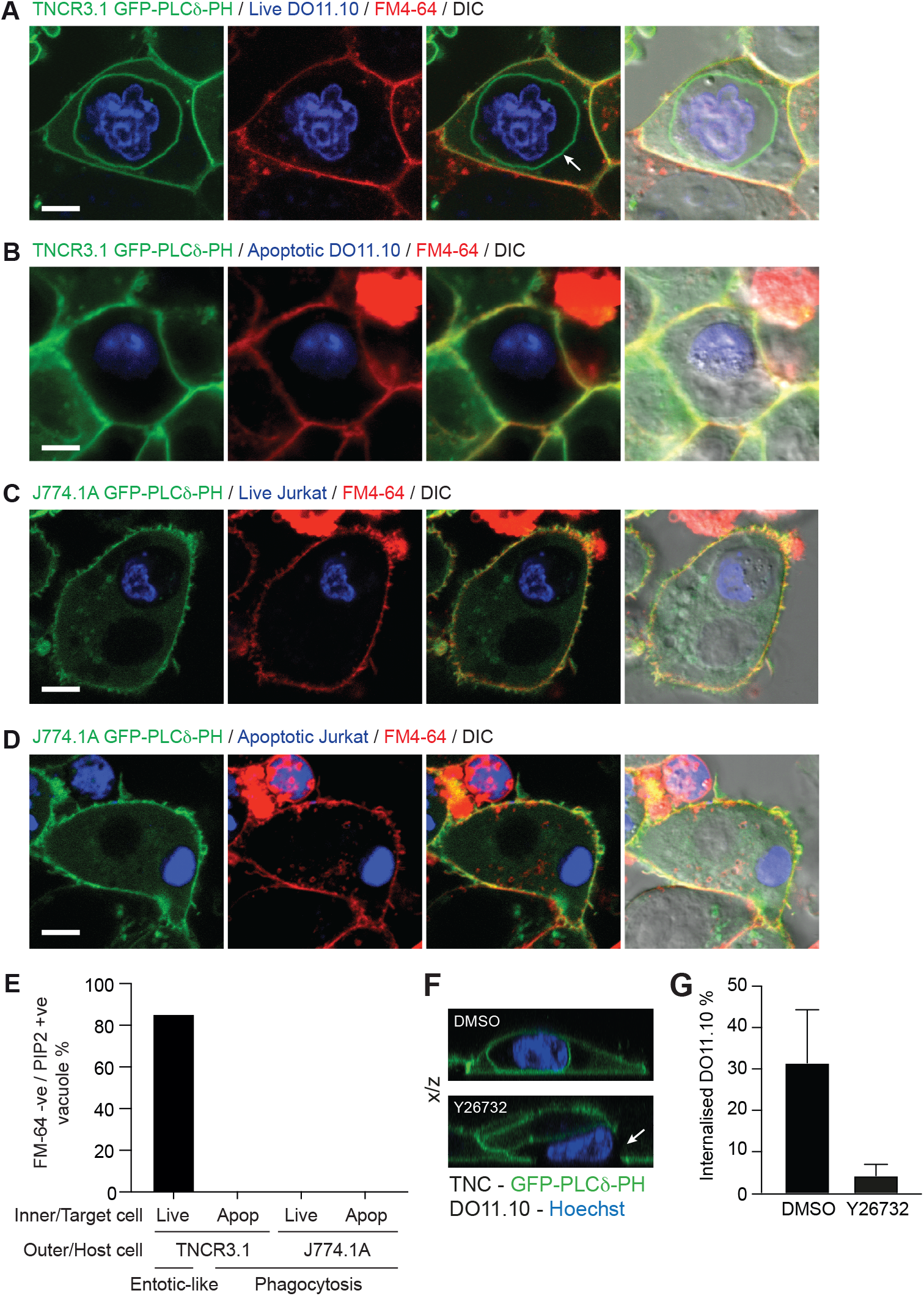
Impact of cargo viability and engulfment mechanism on vacuolar PI(4,5)P2. (A-D) Fluorescent and DIC confocal images of different co-culture interactions, where outer cells express GFP-PLCδ-PH (PIP2, green), inner cells are pre-labelled with Hoechst (DNA, blue) and cultures are exposed to FM4-64 (red). Scale bars, 5 μm. (A) Entotic-like co-culture of TNCR3.1 and live DO11.10 cells. (B) TNCR3.1 phagocytosis of apoptotic DO11.10 cells. (C) J774.1A phagocytosis of live Jurkat cells coated with anti-CD47 antibodies. (D) J774.1A phagocytosis of apoptotic Jurkat cells. (E) Quantification of FM4-64 negative, PI(4,5)P_2_ positive vacuoles/phagosomes in (A-D), n= 50. (F-G) TNCR3.1/DO11.10 cultured as in (A), -/+ Y26732, 10 μM (F) Confocal x/z images; arrow marks incomplete internalisation. (G) Quantification of DO11.10 internalisation; mean±SD from 8 fields of view.

To test this hypothesis further, the mechanistic requirements of TNC/T-cell internalisation were assessed. Entosis is driven by biophysical differences between a pair of epithelial cells, with the stiffer, more contractile cell actively pushing into a softer, more deformable neighbour, dependent on Rho and Rho associated Kinase (ROCK) activity (Overholtzer et al., 2007); by contrast, ROCK does not influence phagocytosis (Olazabal et al., 2002). We found that ROCK inhibition does not impair the T-cells’ ability to migrate under TNCR3.1 cells (Fig. 3F), but it significantly reduces cell-in-cell formation (Fig. 3G), consistent with invasive, entosis-like engulfment.

Together, these data indicate that the internalisation of T-cells by TNCs proceeds through a ROCK-dependent, entosis-like mechanism, and that this mode of engulfment leads to formation of an intracellular, PI(4,5)P_2_-positive vacuole.

### PI(4,5)P_2_ loss precedes, and is required for, inner cell death

Finally, we questioned the functional significance of vacuolar PI(4,5)P2. During phagocytosis, loss of PI(4,5)P_2_ is required for closure and completion of phagosomes (Scott et al., 2005). To investigate the effects of PI(4,5)P2 during entosis-like engulfments, we tracked their longer-term dynamics. During TNC/T-cell internalisation, PI(4,5)P_2_ persisted post-engulfment, as expected. Notably, however, an abrupt loss was then observed over time, occurring prior to the death of target DO11.10 cells, as assessed by DIC morphology and loss of movement (Fig. 4A, Movie 3). These data point to a possible relationship between loss of PI(4,5)P_2_ and cell death.

**Figure 4.**
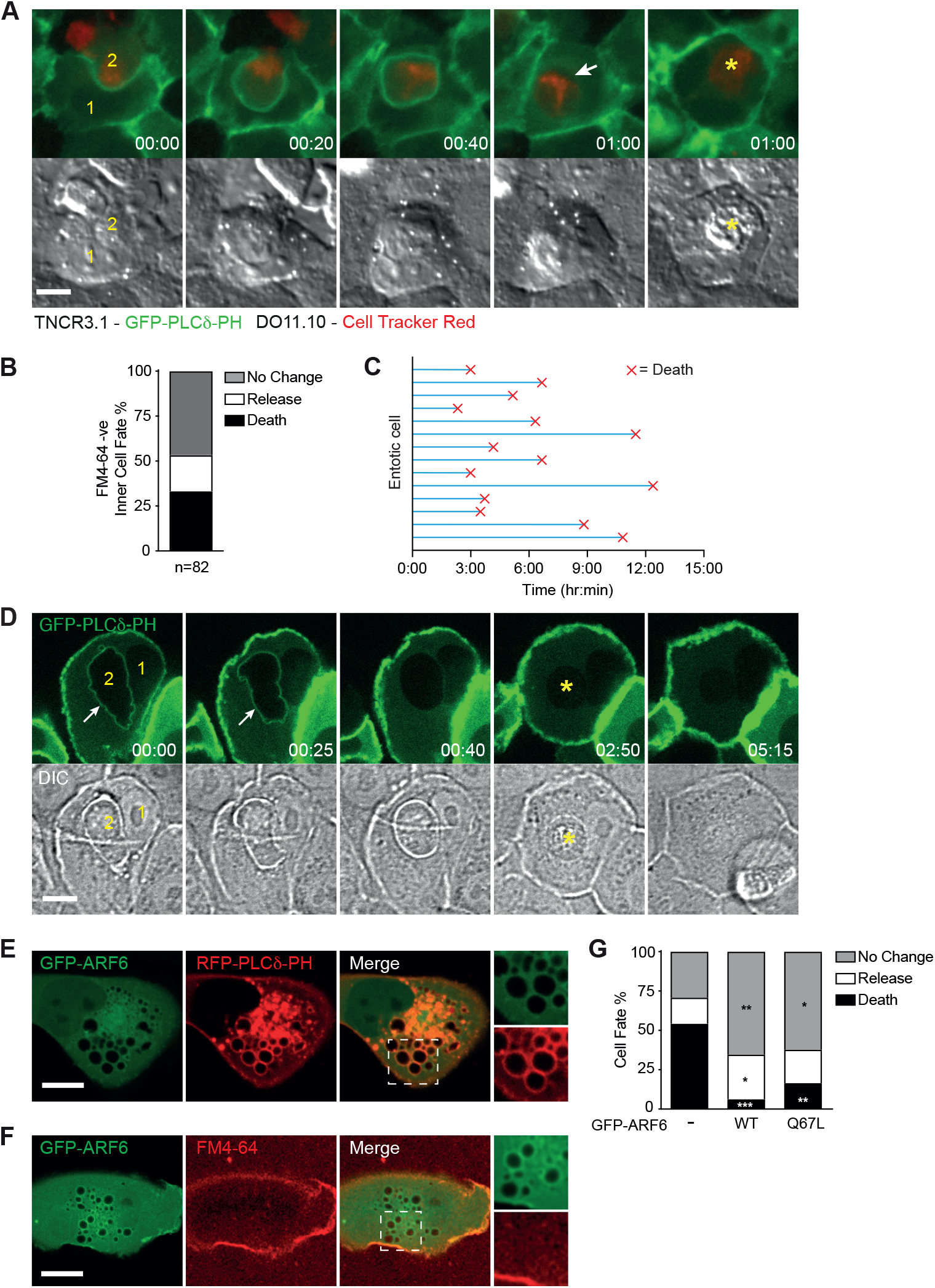
Vacuolar PI(4,5)P2 dynamics influence inner cell death. (A) Wide-field fluorescent and DIC timelapse images of TNCR3.1 cells, expressing GFP-PLCδ-PH (1), interacting with CellTracker Red labelled DO11.10 (2). Arrow marks loss of GFP-PLCδ-PH (PIP2) from vacuole, asterisk marks death of DO11.10 cell. Time, min:sec. Scale bar, 10 μm. (B-D) MCF10A cell entosis, induced by matrix deadhesion. (B) Quantification of inner cell fate in FM4-64 stained cultures, over 20hrs. Data represent means of 3 independent experiments, n=82. (C) Time-course: X marks point of inner cell death in 14 structures from (B). (D) Live confocal timecourse of entosis. Outer cell expressing GFP-PLCδ-PH (1), inner cell (2). Arrow marks GFP-PLCδ-PH (PIP2) on entotic vacuole, asterisk marks death of inner cell. Time, hour:min. Scale bar, 10 μm. (E-F) Representative confocal images of MCF10A cells expressing GFP-ARF6 and (E) RFP-PLCδ-PH (PIP2, red) or (F) FM4-64 (red). Zoomed insets show intracellular vesicles. Scale bars, 5 μm. (G) Quantification of entotic inner cell fate in control MCF10A (n=79), or cells expressing GFP-ARF6 WT (n=137) or Q67L (n=70). Data represent means from 3 independent experiments. *, <p0.02; **, <p0.005; ***, <p0.0002.

To investigate further, we turned to entosis in MCF10A cells, which yields 3 distinct inner cell fates: 1) remains viable within the vacuole; 2) escapes from the host cell; 3) dies, through a non-apoptotic cell death program involving lysosome fusion (Florey et al., 2011) (Fig. 4B). Notably, inner cell death occurred asynchronously over the course of the experiment (Fig. 4C), indicating that formation and maturation of entotic vacuoles are temporally separate events. Moreover, similar to the TNC/T-cell system, loss of PI(4,5)P_2_ consistently occurred prior to inner cell death (Fig. 4D). These data indicate that during entosis, vacuole scission and maturation are uncoupled, and that loss of PI(4,5)P_2_ precedes inner cell death. Based on these findings, alongside the TNC-T-cell data described above, we hypothesized that PI(4,5)P_2_ loss may represent an obligate step in cell killing.

To test this, we modulated PI(4,5)P_2_ levels by manipulating ADP-ribosylation factor 6 (ARF6), which regulates its generation at the plasma membrane through activation of phosphatidylinositol 4-phosphate 5-kinase (PIP 5-kinase) (Honda et al., 1999).

Overexpression of wild type (WT) or constitutively active (Q67L) ARF6 drives accumulation of PI(4,5)P_2_ on endosomal membranes (Brown et al., 2001), due to impeded ARF6 cycling and prolonged recruitment of PIP 5-kinase. Consistently, expression of GFP-ARF6 WT or Q67L in MCF10A cells led to accumulation of PI(4,5)P_2_ on basal intracellular vesicles (Fig. 4E), which were also FM4-64 negative (Fig. 4F). Using this model to ‘offset’ PI(4,5)P_2_ loss during entosis, we found that ARF6 (WT or Q67L) overexpression potently inhibits inner cell death, significantly increasing the proportion of inner cells that remain viable (Fig. 4G). In line with these data, over-expression of GFP-PLCδ-PH, which also interferes with PI(4,5)P_2_ dynamics (Szymańska et al., 2008), similarly reduces entotic cell death (Lee et al., 2019). We conclude that loss of PI(4,5)P_2_ from entotic vacuoles is necessary for their maturation and subsequent killing and cannibalism of inner cells.

In summary, we identify a distinctive, intracellular, PI(4,5)P_2_-positive vacuole formed during entosis and thymic nurse cell/T-cell internalisation. Formation of this compartment depends on a non-phagocytic mode of engulfment, through which one cell actively invades into its host, in a ROCK-dependent fashion. The internalised cell becomes fully engulfed in an endolysosomal vacuole that retains PI(4,5)P_2_, a species usually associated with plasma membrane identity, over an extended period, sustaining its viability. Interestingly, the entotic vacuole can also bear other plasma membrane characteristics, including E-cadherin and EGFR (Florey et al., 2011). Eventual loss of PI(4,5)P_2_ is a vital step in vacuole maturation, essential for inner cell killing by lysosomal degradation.

A key remaining question is what triggers the eventual loss of PI(4,5)P_2_ from entotic-like vacuoles to instigate maturation and cell killing? For phagosomes, PI(4,5)P_2_ regulation is coupled to formation and scission, involving recruitment of lipid phosphatases, loss of kinases and metabolism to Ins(1,4,5)P_3_ and DAG (Bohdanowicz et al., 2012; Fairn et al., 2009; Scott et al., 2005). However, in the context of entosis, vacuole formation and maturation are uncoupled, and further work will be required to define the alternative mechanism. It would also be interesting to investigate the functional significance of this compartment. For instance, in thymic nurse cells, could PI(4,5)P_2_ vacuoles provide a specialised, intracellular niche to ‘shelter’ and support selection of thymocytes/T-cells? Similarly, might the entotic vacuole allow inner cells to evade the immune system?

More broadly, this distinctive PI(4,5)P_2_ -positive compartment, and its processing, influence the duration of vacuole residency, and the fate of the internalised cell. In the context of cancer, regulation of these dynamics would bear important implications for tumour promotion, suppression and evolution through entosis. Vacuole dynamics may also impinge similarly on the functional outcomes of other engulfment events, thereby opening up interesting new questions for future research in this field.

## Acknowledgements

We thank Florey Lab members, Dr Michael Overholtzer and the BI imaging facility. This work was supported by the BBSRC (BB/P013384/1; BBS/E/B/000C0432 and BBS/E/B/000C0434), Cancer Research UK (Career Development award C47718/A16337), MRC/BBSRC/EPSRC (MR/K01580X/1) and the Francis Crick Institute which receives its core funding from Cancer Research UK (CC1076), the UK Medical Research Council (CC1076), and the Wellcome Trust (CC1076).

## Materials and Methods

### Reagents

The following reagents were used: Latrunculin B (L5288; Sigma-Aldrich), EDTA (E6758; Sigma-Aldrich), Y26732 (1254; Tocris), Alexa Fluor 488-phalloidin (8878S; Cell Signalling), Hoechst 33342 (4082S; NEB), FM4-64 (T3166; ThermoFisher), CellTracker Red (ThermoFisher; C34552), IFNγ (315-05; Peprotech).

### Plasmids and constructs

GFP-C1-PLCdelta-PH was a gift from Tobias Meyer (Addgene plasmid # 21179). To generate a retroviral version, GFP-PLCδ-PH was cut out using AgeI/BclI and cloned into pQCXIP (Clonetech) using AgeI/BamHI restriction sites. iRFP-PH-PLCdelta1 was a gift from Pietro De Camilli (Addgene plasmid # 66841). pTK92_Lifeact-GFP was a gift from Iain Cheeseman (Addgene plasmid # 46356). pBabe GFP-PH-Akt was a gift from Dr Len Stephens. pARF6-CFP and pARF6(Q67L)-CFP were gifts from Joel Swanson (Addgene plasmid # 11382 and # 11387). To generate retroviral versions, CFP was replaced with GFP from pEGFP-N1 using AgeI/NotI restriction sites. ARF6-GFP was then cloned into pLPCX (Clonetech) using EcoRI/NotI digests.

### Cell lines and tissue culture

MCF10A cells (human breast epithelial) were cultured in DMEM/F12 (11320074; Gibco) containing 5% horse serum (16050-122; Thermo Fisher Scientific), EGF (20 ng/ml; AF-100-15; PeproTech), hydrocortisone (0.5 mg/ml; H0888; Sigma-Aldrich), cholera toxin (100 ng/ml; C8052; Sigma-Aldrich), insulin (10 μg/ml; I9278; Sigma-Aldrich), and penicillin/streptomycin (100 U/ml; 15140-122; Gibco). J774 mouse macrophages (ATCC) HEK293FT cells (ATCC), and the T cell hybridoma DO11.10 (a kind gift from Dr Philippa Marrack) were cultured in DMEM + 10% FBS with penicillin/streptomycin (100 U/ml; 15140-122; Gibco). Jurkat human T lymphocyte cells (ATCC) were cultured in RPMI-1640 containing 10% FBS (F9665; Sigma-Aldrich) and penicillin/streptomycin (100 U/ml; 15140-122; Gibco). The thymic nurse cell line TNCR3.1 (a kind gift from Dr Willem van Ewijk) were cultured in DMEM containing 10% FBS (F9665; Sigma-Aldrich), penicillin/streptomycin (100 U/ml; 15140-122; Gibco) and 1% β-mercaptoethanol.

### Retrovirus production and infection to generate stable cell lines

HEK293FT cells were transfected with retroviral constructs and retroviral Gag/Pol and VSV-G packaging vectors, using Lipofectamine 2000. Supernatant containing viral particles were collected for 2 days and stored at -80°C. MCF10A, J774.1A or TNCR3.1 cells were seeded in a 6-well plate at 5 × 10^4^ per well. The next day 1 ml viral supernatant was added with 10 µg/ml polybrene (H9268; Sigma-Aldrich) for 24 h followed by a media change. Cells were then selected with appropriate antibiotic.

### Entotic soft-agar microscopy

Plain or biosensor expressing MCF10A cells were trypsinised and mixed 1:1 at a final density of 10^5^ cells/six-well on ultra-low adhesion dishes (Costar) for 2 h. Cells were embedded into growth media + 0.4% low melt agarose (Sigma) and plated overnight onto glass-bottomed dishes (MatTek) precoated with polyhema (Sigma, P3932) to prevent cell adherence. Cell pairs were chosen for imaging using a Confocal Zeiss LSM 780 microscope (Carl Zeiss Ltd).

### Apoptotic and live cell phagocytosis assays

Prior to imaging, J774.1A cells were plated overnight on 35 mm glass-bottomed dishes (MatTek) in the presence of 200 U/ml IFNγ. Opzonised zymosan (OPZ) particles were generated by incubating Zymosan A from Saccharomyces (Z4250; Sigma-Aldrich) in human serum (P2918; Sigma-Aldrich) for 30 min at 37°C. The zymosan was then centrifuged at 5,000 rpm for 5 min and then resuspended in PBS at 10 mg/ml. The solution was passed through a 25-gauge needle using a 1-ml syringe several times to break up aggregates. Cells were mounted on a spinning-disk confocal microscope, comprising a Nikon Ti-E stand, Nikon 60 × 1.45 NA oil immersion lens, Yokogawa CSU-X scanhead, Andor iXon 897 EM-CCD camera, and Andor laser combiner. Where required, FM4-64 5 μg/ml was added for 2 mins prior to image acquisition. All imaging with live cells was performed within incubation chambers at 37°C and 5% CO_2_. Image acquisition and analysis were performed with Andor iQ3 (Andor Technology) and ImageJ.

For apoptotic cell phagocytosis, Hoescht stained DO11.10 or Jurkat cells underwent irradiation in a UV Stratalinker 2000. Cells received two consecutive treatments of 8000mJ UV irradiation and were left to undergo apoptosis for 120 min in the incubator at 37°C, 5% CO^2^. 5 x 10^5^ apoptotic cells were then added to 35mm glass-bottomed dishes containing either TNCR3.1 or IFNγ treated J774.1A cells and incubated at 37oC and 5%CO^2^ for 120 min before image acquisition. For live cell phagocytosis, Jurkat cells were incubated with 10 μg/ml anti-CD47 B6H12 antibody (sc-17320; Santa Cruz) for 30 mins on ice. Cells were washed prior to addition to dishes of IFNγ treated J774.1A cells. Images were acquired as above.

### Entosis time-lapse microscopy

To quantify entotic cell fate, 1.5×10^5^ cells were seeded overnight on 35mm glass-bottomed dishes (MatTek, Ashland, MA) and cell-in-cell structures were imaged the next day. Fluorescence and differential interference contrast (DIC) images were acquired every 4 minutes for 20 hours using a Flash 4.0 v2 sCMOS camera (Hamamatsu, Japan), coupled to a Nikon Ti-E inverted microscope, using a 20 × 0.45 NA objective, performed in an incubation chamber at 37°C, with 5% CO_2_. Image acquisition and analysis was performed with Elements software (Nikon, Japan). Internalized cells were scored for release, death or no change. Cell-in-cell structures chosen for analysis were completely enclosed and shielded from external media at the start of imaging, as assessed by FM4-64 staining. The timing and type of internalized cell death was determined by DIC morphology, cessation of movement or, where present, monitoring nuclear markers.

### 3D CLEM

MCF10A cells expressing GFP-PLCδ-PH were seeded onto 35-mm gridded glass-bottom dishes (# P35G-2-14-CGRD; MatTek) and used the following day. Live-cell images were acquired with a Confocal Zeiss LSM 780 microscope (Carl Zeiss Ltd) at 37°C in a temperature- and CO_2_-controlled chamber, using Zen software (Carl Zeiss Ltd). Cells were fixed by adding 8% formaldehyde (F8775; Sigma-Aldrich) in 0.2 M phosphate buffer pH 7.4, in a 1:1 ratio with culture medium in the dish. Samples were further fixed in 2.5% glutaraldehyde (G5882; Sigma-Aldrich) and 4% formaldehyde in 0.1 M phosphate buffer pH 7.4 for 30 min at room temperature. Samples were then kept in 1% formaldehyde in 0.1 M phosphate buffer pH 7.4 and stored at 4°C until processing. For Serial Block Face Scanning Electron Microscopy (SBF-SEM), cells were prepared and mounted as described previously (Russell et al., 2017). In brief, blocks were trimmed to a small trapezoid, excised from the resin block, and attached to SBF-SEM specimen holders using conductive epoxy resin. Prior to commencement of the SBF-SEM imaging run, the samples were coated with a 2nm layer of platinum to further enhance conductivity. SBF-SEM data was collected using a 3View2XP (Gatan, Pleasanton, CA) attached to a Sigma VP SEM (Carl Zeiss Ltd, Cambridge, UK). Inverted backscattered electron images were acquired through the entire volume of each cell of interest. For each 50 nm slice, a high-resolution image of the cell was acquired with the following parameters: 8192 x 8192 pixels, 2 µs dwell time and pixel size of 5.2 nm (Fig. 1J) and 7.8nm (Fig. S1). The SEM was operated in variable pressure mode, at an accelerating voltage of 2 kV with a 20 µm aperture.

### Statistical analysis

Statistical analysis was performed using GraphPad Prism software. One-way ANOVA followed by Tukey multiple comparison test was used as indicated in figure legends.

### Schematics

All schematics were generated using Biorender.

