## Supplementary Material for "A distinctive PI(4,5)P_2_ compartment forms during entosis and related engulfment processes"

**Figure S1**

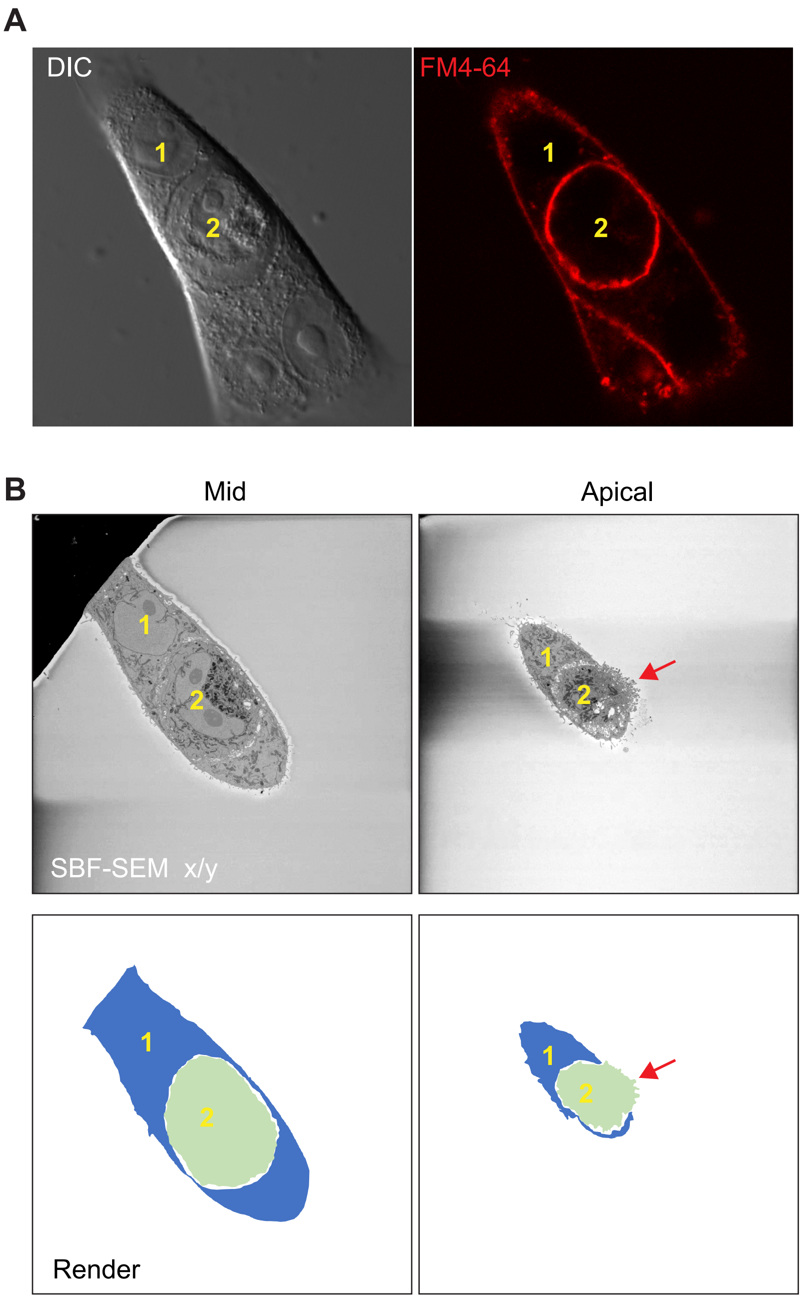


3D CLEM analysis of partial MCF10A entotic structure, outer cell (1), inner cell (2). (A) Live cell confocal microscopy images of DIC and FM4-64. (B) SBF-SEM x/y slice and cartoon render. Arrow denotes inner cell protruding out of host.

**Movie 1**

RAW264.7 cell expressing PI(4,5)$P_2$ sensor GFP-PLCδ-PH, phagocytosing opsonized zymosan particle (asterisk). Time min:sec.

**Movie 2**

SBF-SEM 3D CLEM x/y axis stack of MCF10A entotic cell-in-cell structure, related to Figure 1J.

**Movie 3**

Fluorescent and DIC time-lapse of CellTracker Red labelled DO11.10 cells interacting with GFP-PLCδ-PH expressing TNCR3.1cells. Movie shows DO11.10 cell entering TNCR3.1 forming a PI(4,5)$P_2$ positive vacuole. PI(4,5)$P_2$ is lost prior to the death of inner cell. Related to Figure 4 A.
